# Interdependence of a microtubule polymerase and a motor protein in establishment of kinetochore end-on attachments

**DOI:** 10.1101/2023.06.08.544255

**Authors:** Julia R. Torvi, Jonathan Wong, David G. Drubin, Georjana Barnes

## Abstract

Faithful segregation of chromosomes into daughter cells during mitosis requires formation of attachments between kinetochores and mitotic spindle microtubules. Chromosome alignment on the mitotic spindle, also referred to as congression, is facilitated by translocation of side-bound chromosomes along the microtubule surface, which allows the establishment of end-on attachment of kinetochores to microtubule plus ends. Spatial and temporal constraints hinder observation of these events in live cells. Therefore, we used our previously developed reconstitution assay to observe dynamics of kinetochores, the yeast kinesin-8, Kip3, and the microtubule polymerase, Stu2, in lysates prepared from metaphase-arrested budding yeast, *Saccharomyces cerevisiae*. Using total internal reflection fluorescence (TIRF) microscopy to observe kinetochore translocation on the lateral microtubule surface toward the microtubule plus end, motility was shown to be dependent on both Kip3, as we reported previously, and Stu2. These proteins were shown to have distinct dynamics on the microtubule. Kip3 is highly processive and moves faster than the kinetochore. Stu2 tracks both growing and shrinking microtubule ends but also colocalizes with moving lattice-bound kinetochores. In cells, we observed that both Kip3 and Stu2 are important for establishing chromosome biorientation, Moreover, when both proteins are absent, biorientation is completely defective. All cells lacking both Kip3 and Stu2 had declustered kinetochores and about half also had at least one unattached kinetochore. Our evidence argues that despite differences in their dynamics, Kip3 and Stu2 share roles in chromosome congression to facilitate proper kinetochore-microtubule attachment.

## Introduction

Mitosis is an essential process by which replicated chromosomes are segregated to daughter cells. To achieve this, chromosomes are physically partitioned into daughter cells via a microtubule-mediated process. Through a complex process involving microtubule assembly and disassembly, combined with actions of kinesin and dynein motor proteins, forces first align the chromosomes on the metaphase plate and then separate the sister chromosomes to the two spindle poles. Spindle microtubules attach to a specialized protein structure assembled on centromere DNA called the kinetochore. The kinetochore binds both to the sides and plus ends of microtubules and can move along microtubules via lateral interactions with the microtubule lattice. It also can move in association with the microtubule plus end and allows tubulin association and dissociation with the plus end while remaining tightly bound (Biggins et al., 2013).

The process by which chromosomes align at the center of the mitotic spindle for proper partitioning is called chromosome congression (Maiato et al., 2017). Chromosome congression is a very complex process dependent upon motor forces (Cottingham et al., 1999), spindle geometry (Risteski et al., 2021), post-translational modifications (Barisic and Maiato, 2016), and chemical gradients (Heald and Khodjakov, 2015). Many studies have addressed the roles of motor proteins in chromosome congression, especially in mammalian cells. Dynein, a minus-end directed microtubule motor protein, pulls chromosomes along astral microtubules toward the centrosome, facilitating attachment of kinetochores to microtubule plus ends (Kapoor et al., 2006; Li et al., 2007). Chromosomes that do not make end-on, plus end, attachment to kinetochore-microtubules can be transported to the plus-end by a kinesin-7, CENP-E (Craske and Welburn, 2020; Lemura and Tanaka, 2015). When CENP-E is chemically inhibited in cells, kinetochores do not align on the metaphase plate. Instead, many chromosomes remain near the poles, with their distribution spanning the entire spindle length, resulting in mitotic arrest and even cell death (Barisic et al., 2015).

In budding yeast, several proteins have been shown to provide functions similar to those implicated in facilitating chromosome congression in mammals. The yeast minus end-directed kinesin-14, Kar3, is functionally analogous to dynein, moving chromosomes toward the spindle pole body (Tanaka et al., 2005). Our previous research provides evidence that the yeast kinesin-8, Kip3, has a function similar to CENP-E. When Kip3 is absent in cells, kinetochores are spread along the mitotic spindle rather than aligned along the metaphase plate (Torvi et al., 2022). Additionally, Kip3 absence from *in vitro* assays results in loss of plus end-directed kinetochore motility (Torvi et al., 2022). However, the molecular mechanism by which Kip3 mediates kinetochore motility remains a mystery. Kinetochores move along microtubules much more slowly than Kip3 itself. Additionally, kinetochores frequently pause on the microtubule lattice, whereas Kip3 movement is constant. Therefore, Kip3-mediated kinetochore transport along microtubules appears distinct from behavior observed for transport of other cargos along microtubules by motor proteins. Here we set out to further explore Kip3’s involvement in kinetochore motility.

One candidate for a protein that might mediate Kip3’s role in kinetochore motility is the microtubule polymerase, Stu2. Split fluorescent protein assays, yeast two-hybrid screens, and immune-purifications indicate that Kip3 and Stu2 interact physically (Gandhi et al., 2011). In addition to Stu2’s microtubule polymerase function, Stu2 has recently been shown to have a role at the kinetochore facilitating proper microtubule attachments. Through immune-purifications, binding assays, cross-linking mass spectrometry, and crystallography, Stu2 has been shown to bind to kinetochores via interaction with the Ndc80 complex and aide in the establishment of proper kinetochore-microtubule attachments (Miller et al., 2016, 2019; Zahm et al., 2021). However, we recently showed that, unexpectedly, Stu2’s essential function is not its microtubule polymerase activity, but rather an as-yet-to-be defined function that requires nuclear localization (Carrier et al., 2022). Given that Kip3 binds to Stu2, and that Stu2 can bind to the kinetochore, we set out to explore possible functional interdependence between Kip3 and Stu2 in kinetochore motility during congression. Using the assay we developed previously to observe single kinetochores moving on microtubules, and to implicate Kip3 in this process, we now investigated possible Kip3-Stu2 functional cooperation during kinetochore motility (Torvi et al., 2022; Bergman et al., 2018).

## Results

### Stu2 is required for kinetochore movement toward the microtubule plus end

Previously we reported that kinetochores bind to microtubules in metaphase-arrested cell lysates and translocate on the lateral surface toward the plus end (Torvi et al., 2022). While kinesin-8, Kip3, absence from lysates resulted in immobile kinetochores, the mechanism of lateral directional kinetochore motility was, nevertheless, not clear since the kinetochore movement rate was markedly slower than Kip3’s velocity. Here, we further investigated this mechanism by determining if Stu2, a protein that binds both to kinetochores and to Kip3, is involved in this process. Briefly, using our reconstitution assay (Bergman et al., 2018; Torvi et al., 2022), which uses *Saccharomyces cerevisiae* cell-cycle arrested whole cell lysates, we observed dynamic kinetochores and MAPs on microtubules via TIRF microscopy. Kinetochore proteins (Spc105) and MAPs (Kip3, Stu2) were expressed at endogenous levels as fusions to fluorescent proteins in a strain background with fluorescently tagged alpha-tubulin (Tub1) and a temperature sensitive mutant (*cdc23-1*) that causes cells to arrest in metaphase at the non-permissive temperature (Irniger et al., 1995).

As we reported previously, in wild-type lysates, kinetochores (tracked by imaging Spc105-GFP), bind to the lateral surface of microtubules and move in a processive, directional manner toward the microtubule plus end at a speed of 0.57±0.041 μm/min and average run length of 0.81±0.059 μm (Torvi et al., 2022) (Figure 1A-D, Movie 1). When bound to the lattice, the kinetochores spend most of the time stationary, but spend 36% of the time moving toward the plus-end and <1% of the time moving toward the minus-end (Figure 1B). Due to both kinetochore movement toward the plus end and microtubule plus end dynamic assembly and disassembly (Bergman et al., 2018), kinetochores eventually reach microtubule plus-ends and establish end-on attachments. Once a kinetochore is bound to the microtubule plus end, that end is never observed to resume growth. Instead, kinetochore-bound plus ends are either paused (81% of the time) or shrinking (19%) (Torvi et al., 2022) (Figure 1, A and B).

**Figure 1.**
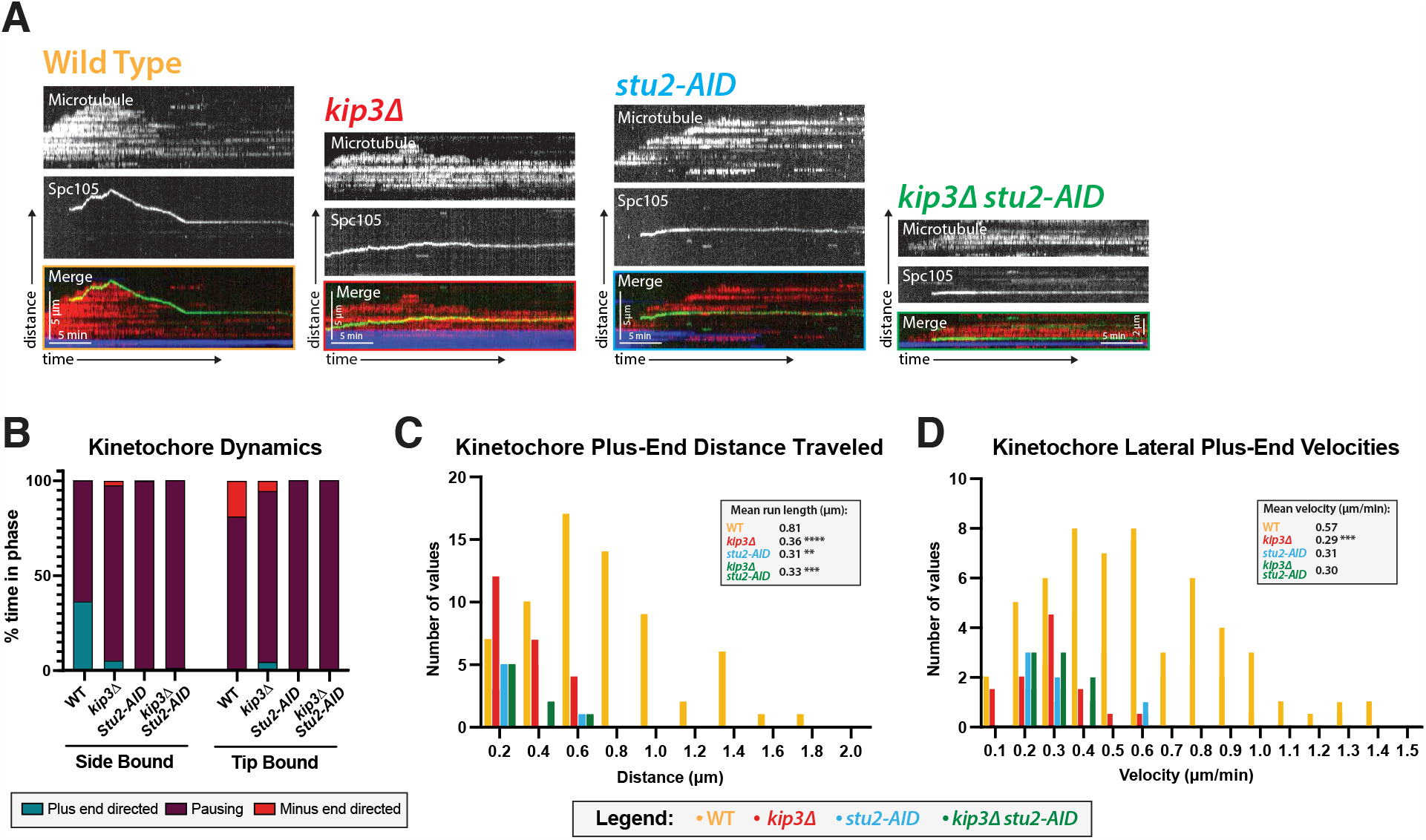
Kip3 and Stu2 are required for lateral kinetochore movement on metaphase microtubules. (**A**) Kinetochore movement toward the microtubule plus end is dependent on Kip3 and Stu2. Representative kymographs from metaphase-arrested lysates with Spc105-GFP, an outer kinetochore protein. Time is on the x-axis and distance is on the y-axis (scale bars are 5 min and 5 μm, respectively). The kinetochore (Spc105-GFP) is shown in green, the yeast microtubule (mRuby2-Tub1) is red, and the stabilized microtubule seed is blue. In wild-type lysates kinetochores toward the microtubule plus-end when bound to the microtubule lateral surface and track the microtubule tip after becoming end-bound. (**B**) Quantification of dynamics over time shows that kinetochores spend 36.16% of their time moving toward the plus-end of the microtubule when laterally bound to the microtubule. This value decreases to 5.18%, 1.13%, and 1% in *kip3*Δ, *stu2-AID*, and *kip3*Δ *stu2-AID* lysate, respectively. Additionally, when a kinetochore reaches the microtubule end in a mutant lysate, reduced tip-bound, minus-end directed kinetochore movement is observed compared to wild-type lysate. (**C**) Both run lengths and the number of runs the kinetochores make toward the microtubule plus-ends depend on Kip3 and Stu2. While the wild-type run length was 0.81 μm in *kip3*Δ, *stu2-AID*, and *kip3*Δ *stu2-AID* lysate lysates showed run lengths reduced to 0.36 μm, 0.31 μm, and 0.33 μm, respectively. Statistical analysis was done by a Kruskal-Wallis test, wherein **** is p <0.0001, *** is p = 0.0005, and ** is p = 0.0021. (**D**) Laterally bound kinetochore velocities were reduced from 0.57 μm/min to 0.29 μm/min, 0.31 μm/min, and 0.30 μm/min in *kip3*Δ, *stu2-AID* and *kip3*Δ *stu2-AID* lysates, respectively. Statistical analysis was done using a Kruskal-Wallis test where *** is p=0.0004. (**B-D**) Quantification is from two replicate trails. For each strain from WT to *kip3*Δ *stu2-AID* lysate in the order listed, N=42, 47, 38, and 46 Spc105-GFP proteins tracked.

So far, based on our previous observations of mutants of all yeast kinesins, we found that only the kinesin-8 Kip3 (Torvi et al., 2022) is necessary for motility. Here, we found that the microtubule polymerase, Stu2 (Figure 1, Movie 2, Movie 3, Movie 4), is also necessary for the directional and processive movement toward the plus end of the kinetochore bound to the microtubule lattice. When Kip3, Stu2, or both of these proteins, is absent, motility is defective (Figure 1A). In the three mutant lysates: *kip3*Δ, *stu2-AID*, and *kip3*Δ *stu2-AID* mutant, the kinetochore run lengths decreased to 0.36±0.28, 0.31±0.040, and 0.33±0.045 μm, respectively (Figure 1C).

Similarly, the kinetochore velocities decreased to 0.29±0.028, 0.31±0.051, and 0.30±0.031 μm/min, respectively. However, only when Stu2 was absent was the decreased velocity statistically significant (Figure 1D). Lastly, we observed altered microtubule dynamics when Kip3 and Stu2 were removed. Both proteins have known microtubule dynamics functions, which have been characterized previously (Su et al., 2011; Arellano-Santoyo et al., 2017; Ayaz et al., 2012) and also analyzed using this assay (Bergman et al., 2018; Carrier et al., 2022). Kip3 is a catastrophe factor, so microtubules depolymerize less in the *kip3*Δ (Bergman et al., 2018). Stu2 is microtubule polymerase and also contributes to microtubule catastrophe, so we also observe microtubules in a *stu2-AID* growing slowly and rarely depolymerizing (Carrier et al., 2022). Interestingly, the loss of Kip3 and Stu2, individually or together, also abolishes tip-bound kinetochores moving with shrinking microtubules, implying that these MAPs are necessary for both lateral kinetochore movement and kinetochore-induced microtubule depolymerization (Figure 1B).

### Distinct Kip3 and Stu2 dynamics on microtubules in metaphase lysates

Given that the absence of either Kip3 or Stu2 alone, or in combination, results in kinetochores that no longer move processively on the microtubule lattice, we next characterized the dynamics of these proteins in metaphase-arrested lysates to look for evidence of a functional relationship. As shown previously (Bergman et al., 2018; Torvi et al., 2022), Kip3 is highly abundant on microtubules and walks processively toward the microtubule plus-end at a velocity of 2.8±0.047 μm/min without pausing (Figure 2A, Movie 5). Additionally, the fluorescence intensity of Kip3 on the growing microtubule end was about twice as high as on the shrinking microtubule end (Figure 2A). This dynamic behavior in metaphase arrested lysates is distinct from the behavior observed in lysates of cells arrested in other cell cycle stages (Supplement Figure 1A). In S-phase and anaphase lysates, Kip3 was not processive and came on and off the microtubule dynamically. In conclusion, Kip3 processivity and plus-end accumulation were specific to metaphase-arrested lysates.

**Figure 2.**
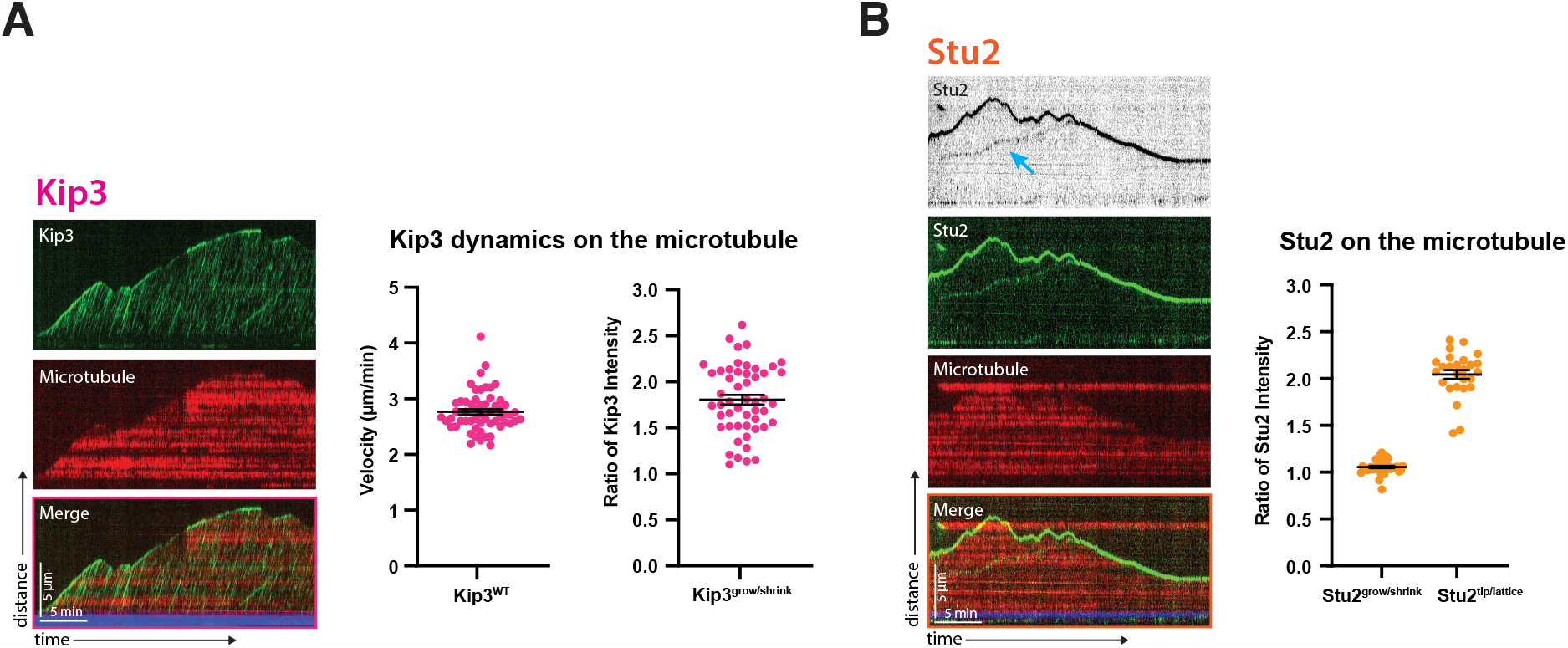
Distinct Kip3 and Stu2 dynamics on the metaphase microtubule. **(A)** Representative kymograph showing Kip3-GFP in green, the yeast microtubule (mRub2-Tub1) in red, and the porcine seed in blue in a metaphase-arrested lysate. Time is on the x-axis and distance is on the y-axis (scale bars are 5 min and 5 μm, respectively). Processive Kip3 tracks are seen streaming along the microtubule with Kip3 accumulation at the plus ends of growing microtubules. The mean velocity of Kip3 is 2.8 μm/min and the ratio of Kip3 fluorescent intensity on growing microtubule ends compared to shrinking ends is 1.8. On both graphs the SEM is indicated in black. **(B)** Representative kymograph showing Stu2-GFP in green, the yeast microtubule (mRuby2-Tub1) in red, and the porcine seed in blue in a metaphase-arrested lysate. Time is on the x-axis and distance is on the y-axis (scale bars are 5 min and 5 μm, respectively). Stu2 tracks both the growing and shrinking ends of microtubules. Additionally, a dimmer Stu2 signal is apparent on the microtubule lattice, slowly moving toward the tip in a unidirectional manner (blue arrow). This localization is quantified in the graph to the right, where the ratio of Stu2 fluorescence intensity at growing and shrinking microtubule ends is 1.05 and the ratio at the tip compared to the lattice is 2.05. The SEM is indicated in black.

The dynamics of Stu2 were distinct from those of Kip3. As previously reported, we observed Stu2 associated with the microtubule tip, binding tubulin dimers and facilitating their addition to the growing end (Geyer et al., 2018). Interestingly, we also observed that the amount of Stu2 on the tip of polymerizing versus depolymerizing microtubules was equal and steady over time (Figure 2B, Movie 6), a novel characterization not yet observed. This characteristic is notably different from other plus-tip associated proteins like Bim1/EB1 in *in vitro* assays (Vaughan, 2005) since those proteins predominantly associate with growing microtubule ends. In addition to observing Stu2 on microtubule plus ends, we observed a fainter Stu2 signal moving slowly along the microtubule lattice towards the plus-end. Compared to the fluorescent intensity of Stu2 at the microtubule tip, the lattice-bound Stu2 signal was about half as bright (Figure 2B). Another difference when compared to Kip3 was that Stu2 dynamics were unchanged throughout the cell cycle. In all cell cycle stages tested, Stu2 remained on both growing and shrinking microtubule ends, and there was frequently a dimmer lattice-bound signal (Supplement Figure1B). However, plus-end-directed movement of Stu2 along microtubules was only observed in metaphase-arrested lysates. For this reason, we focused our investigation of the roles of Kip3 and Stu2 in metaphase.

### Stu2 co-localizes with kinetochores *in vitro* and acts with Kip3 to facilitate lateral kinetochore movement

We hypothesized that Kip3, Stu2, and kinetochores might associate to facilitate kinetochore motility. To test this possibility, we imaged Kip3-GFP and Stu2-GFP in combination with a fluorescently tagged kinetochore protein, Spc105-mScarlet, and looked for co-localization in lysates made from cells arrested at metaphase. Based on the dynamics of the faint Stu2 signal we observed on lateral microtubule surfaces (Figure 2B), we hypothesized that this signal might represent a kinetochore-bound pool. When visualizing Stu2-GFP in combination with Spc105-mScarlet, we observed that, invariably, the faint lattice-bound Stu2 colocalized with the Spc105 (Figure 3A, Movie 7). In sharp contrast, as reported previously (Torvi et al., 2022), Kip3 moves much faster than the kinetochore and no Kip3 signal was ever detected moving along with the kinetochore, although there is frequent overlap of the kinesin’s tracks with those of Spc105 (Figure 3C, Movie 9).

**Figure 3.**
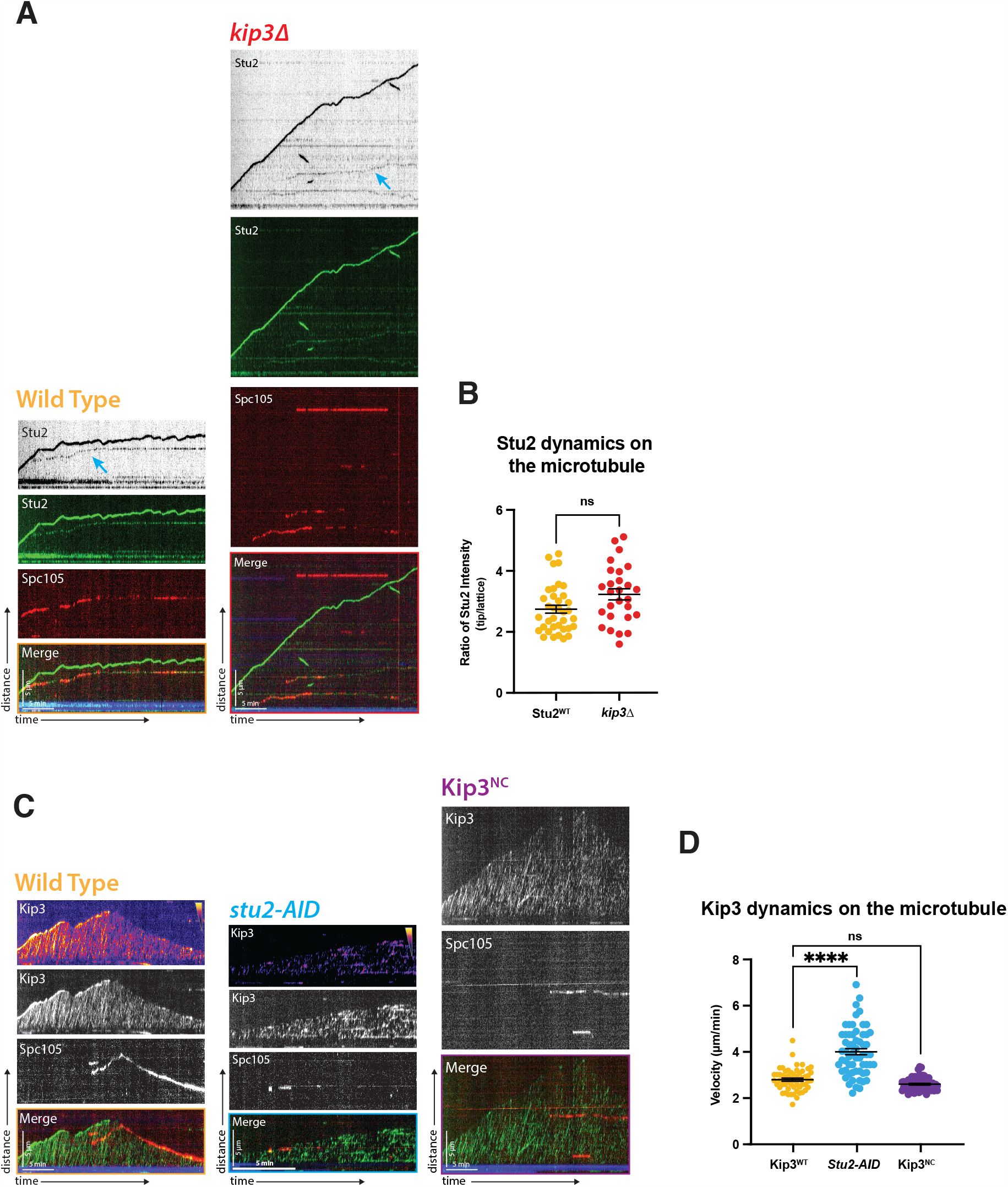
Kinetochore movement toward microtubule plus end depends on Kip3 and association with Stu2. **(A)** Representative kymograph showing metaphase arrested lysate with Stu2-GFP in green, Spc105-mScarlet in red, and the porcine seed in blue. Time is on the x-axis and distance is on the y-axis (scale bars are 5 min and 5 μm, respectively). In wild-type lysate one population of Stu2 tracks the microtubule tip and the other colocalizes with kinetochores. In a *kip3*Δ lysate, Stu2’s association with kinetochores is independent of Kip3. **(B)** Fluorescence intensity of Stu2 at the microtubule tip is 2.8 fold greater than Stu2 associated with kinetochores on the microtubule lattice in wild-type lysates. This ratio is essentially unchanged when Kip3 is absent, with only a statistically insignificant increase to 3.2. The SEM is shown in black. Statistical analysis was done using a Kruskal-Wallis test where ns is p=0.1145. Quantification is from two replicate trials. For lysates from each strain, WT and *stu2-AID*, N=36 and 27, number of Stu2-GFP signals quantified. **(C)** Representative kymograph from lysate made from metaphase-arrested cells with Kip3-GFP in green, Spc105-mScarlet in red, and the porcine seed in blue. Time is on the x-axis and distance is on the y-axis (scale bars are 5 min and 5 μm, respectively). In wild-type lysate, Kip3 moves faster than the kinetochore on the microtubule lattice with no detectable colocalization. When Stu2 is depleted, the kinetochore does not move, but Kip3 moves faster with shorter run lengths. When the Kip3 neck chimera variant replaces the wild-type protein, microtubule-associated kinetochores do not move, but Kip3 is still processive, though it no longer accumulates on growing microtubule ends. **(D)** Kip3 velocities were quantified revealing that Kip3 moves faster upon Stu2 depletion, showing an increase in velocity from 2.8 μm/min to 4.0 μm/min. Additionally, Kip3^NC^ did not show altered speed relative to the wild-type protein, with a velocity of 2.6 μm/min. The SEM is shown in black. Statistical analysis was done using a Kruskal-Wallis test where **** is p<0.0001. Quantification is from two replicate trials. For each strain in the order listed from WT to Kip3^NC^ lysate, N=57, 61, and 64 Kip3 proteins were tracked.

Although no colocalization between Stu2 and Kip3 was detected, we wished to further test the possibility that Stu2 acts as a bridge between Kip3 and the kinetochore since these two proteins have been reported previously to interact with each other. To do this, we tested whether Stu2 localization at kinetochores was dependent on Kip3. When the gene encoding Kip3 is deleted, kinetochores do not move on the microtubule (Figure 1) and microtubules undergo much less frequent catastrophes (Bergman et al., 2018). However, Stu2 colocalization with kinetochores was not detectably altered in this mutant. In the absence of Kip3, Stu2 is still associated with both the growing and shrinking microtubule ends and still colocalizes with kinetochores. Also, the ratio of the intensity of Stu2-GFP at microtubule ends to the intensity at the kinetochore was not detectably different between wild-type and *kip3*Δ lysates, although TIRF is an imperfect approach for making such ratio-metric determinations (Figure 3B, Movie 8).

Given that Stu2 is still at the kinetochore when Kip3 is absent (Figure 3A) and that kinetochores do not move when Stu2 is absent (Figure 1), we hypothesized that Stu2 connects Kip3 to the kinetochore. We predicted that, if we were to observe Kip3 and the kinetochore in the absence of Stu2, Kip3 would show wild-type dynamics. Surprisingly, this was not the case. When Stu2 is absent, as reported above, kinetochores become immobile (Figure 1), microtubule growth slows and depolymerization decreases (Carrier et al., 2022). However, Kip3 dynamics were also severely altered (Figure 3C, Movie 10). Instead of non-stop runs at a velocity of 2.8±0.064 μm/min, runs become very short and increase in velocity to 4±0.13 μm/min in the absence of Stu2 (Figure 3D; note the 2x acquisition rate of the movies in Figure 3C required because of the faster Kip3 motility rates). Additionally, there was less Kip3 bound to microtubules when Stu2 was absent, as shown by the heatmap of Kip3 in kymographs (Figure 3C). Finally, there was a pronounced diminution of Kip3 accumulation at microtubule plus ends in this mutant. Additionally, we were able to show that the addition of wild-type Stu2 lysate back to a *stu2-AID* lysate restored Kip3 processivity (Supplement Figure2A). Reversing this sequence, when *stu2-AID* lysate was replaced by a lysate with wild-type Stu2, Kip3 processivity was instantly diminished (Supplement Figure2A). These results indicate that Stu2 absence affects more than just kinetochore motility along microtubules in this lysate assay. While kinetochores are immobile in a Stu2-depleted lysate, this could be because either it mediates the interaction between Kip3 and the kinetochore or that it affects Kip3 motility, which then results in the loss of kinetochore motility (see Discussion).

As another approach to exploring the functional relationship between Stu2 and Kip3, we next attempted to create a Kip3 separation-of-function mutant. Given that we previously found that kinetochores were still motile in a Kip3 mutant lacking the entire tail (amino acids 482-805) (Torvi et al., 2022), we turned our attention to the 42 amino acid neck-linker (amino acids 439-481). Based on earlier reports that Kip3 amino acids 1-446 can bind to Stu2 (Gandhi et al., 2011), we narrowed our target to amino acids 439-446. Kinesin neck linker sequences are very important for dictating the motor’s processivity (Kim et al., 2014; Malaby et al., 2019). We therefore did not want to mutate the neck in a way that would disrupt its structure. We substituted amino acids 439-446 with those from another similarly processive kinesin, Kif18a, a mammalian kinesin-8 that shares many similarities to Kip3, both in structure and function (Stumpff et al., 2008, 2011). Therefore, we created a Kip3-Kif18a neck-chimera (Kip3^NC^) replacing amino acids 439-446 in Kip3 with those of Kif18a (Supplement Figure3A). Mutating these six amino acids in the neck of Kip3 resulted in a motor with wild-type processivity (Figure 3, C and D), but resulted in immobile kinetochores (Figure 3C, Movie 11). Interestingly, this Kip3^NC^ protein no longer accumulated on the plus-ends of microtubules. These microtubules also exhibited a lower catastrophe frequency, consistent with loss of a previously reported Kip3 activity (see Discussion). Despite this unintended plus end-associated phenotype and our current inability to determine whether this mutant can bind to Stu2, we can conclude that these six amino acids in the neck-linker are necessary for kinetochore motility.

### Kip3 and Stu2 work together for proper metaphase spindle and kinetochore organization

Although the reconstitution experiments described above make possible determining the behavior of different proteins on single yeast microtubules in a cell-cycle specific manner, in the mitotic spindle, microtubule geometry and spindle forces are expected to influence protein behaviors (Trupinić et al., 2022) Therefore, we tested how Kip3 and Stu2 affect kinetochore biorientation in intact spindles within live cells and compared these phenotypes to those detected in our reconstitution assay. As we reported previously, loss of Kip3 results in an increased frequency of kinetochores being declustered along the spindle axis (Torvi et al., 2022) (Figure 4, A and B). We hypothesized that this declustered phenotype is the result of an increased frequency of laterally bound kinetochores that fail to congress to the microtubule plus-end (Figure 1). We now report observing declustered kinetochores in cells expressing Kip3^NC^, indicating that this phenotype is the result of immobile kinetochores and not the result of defective Kip3 motility (Figure 4, A and B). While Stu2 depletion in our assay resulted in immobile kinetochores, Stu2’s role as a microtubule polymerase also leads to very short, slowly growing microtubules. This microtubule phenotype is also seen in mitotic spindles of stu2 mutants (Miller et al., 2016) (Figure 4B). However, in the absence of Stu2, the frequency of unattached kinetochores increased, as reflected in appearance of kinetochore spots far off-axis with the spindle. Additionally, these cells often show both a declustered kinetochore phenotype (more than two kinetochore puncta on axis with the microtubules) and the unattached kinetochore phenotype (Figure 4, A and B). Together, these phenotypes both affirm Stu2’s role in kinetochore attachment (Miller et al., 2016) and suggest a novel role in chromosome congression (this study). Given that loss of Kip3 or Stu2 results in both immobile kinetochores and kinetochore biorientation defects, we sought to gain further insights into how these two proteins work together to facilitate formation of proper kinetochore-microtubule attachments. While the dynamics of these two proteins on microtubules are distinct, and their loss results in different spindle phenotypes, they share a common role in kinetochore congression and biorientation. Therefore, we hypothesized that the simultaneous loss of the two proteins might result in a more dramatic biorientation and spindle defect than either single mutant alone. Strikingly, *kip3*Δ *stu2-AID* cells completely unable to form bilobed kinetochore puncta. All cells had declustered kinetochores with about half also having at least one unattached kinetochore (Figure 4, A and B). This result supports the conclusion (Figure 5) that these two proteins share a role in facilitating kinetochore congression and biorientation.

**Figure 4.**
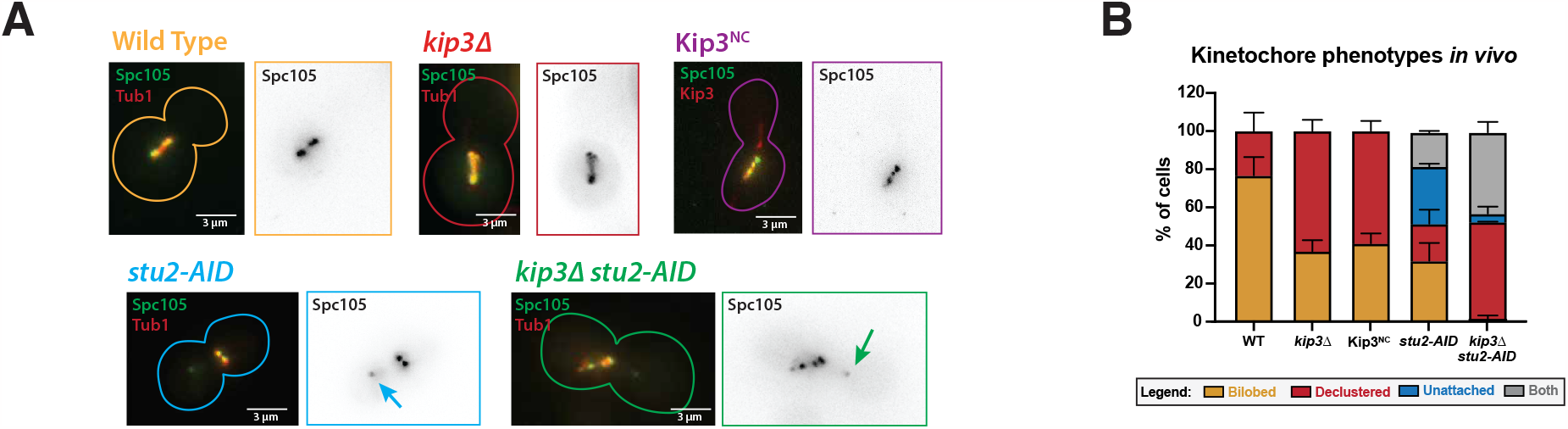
Kip3 and Stu2 work together to achieve proper metaphase spindle and kinetochore organization. **(A)** Metaphase cells with either wild-type KIP3 and STU2 alleles, or *kip3*Δ, *Kip3*^*NC*^, *stu2-AID*, and kip3 and *stu2-AID* alleles, were imaged in a *cdc23-1* strain expressing Spc105-GFP (green) and either mRuby2-Tub1 (red). Note that these cells were not shifted to non-permissive temperature and were not arrested in the cell cycle for imaging because the bud morphology indicates the cell cycle stage. Each image shown is a maximum Z-projection of a 5 μm stack of 0.2 μm slices. **(B)** Quantification of the declustered and unattached kinetochore phenotype is presented as a bar graph for spindles that are 2–3 μm in length. Cells were classified as ‘bilobed’ by a line scan showing two distinct peaks (represented in the wild-type image). The ‘declustered’ phenotype included all line scans that did not show two distinct peaks (represented in the *kip3*Δ image). The ‘unattached’ phenotype was classified as the presence of at least one kinetochore signal off axis away from the main spindle (represented in the *stu2-AID* image). The ‘both’ classification represents spindles that were both declustered and that had an unattached kinetochore (represented in the *kip3*Δ and *stu2-AID* image). Two biological replicates were analyzed, each with n=50 cells. Error bars are standard errors of the mean.

**Figure 5.**
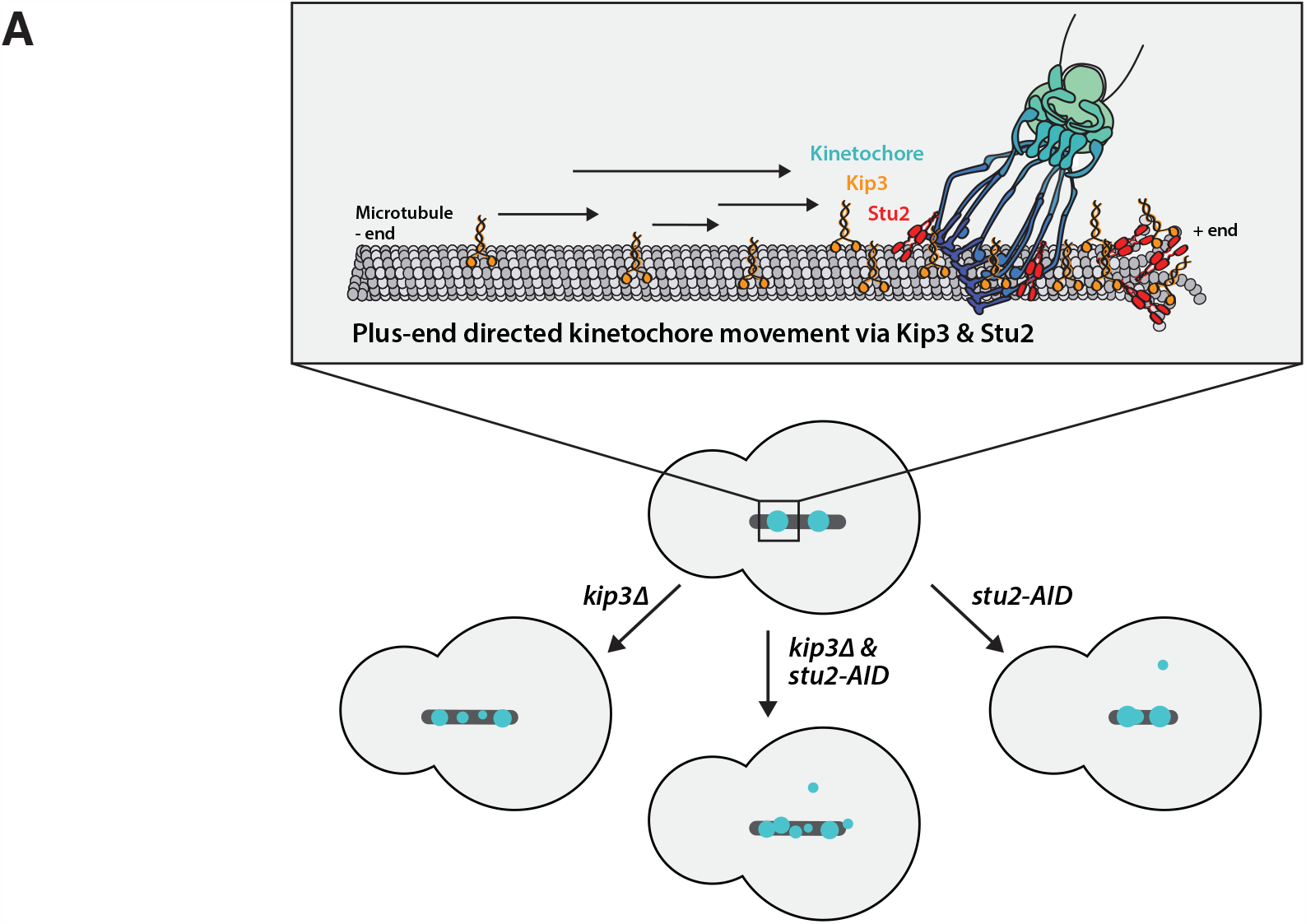
Model for Kip3 and Stu2 mediated kinetochore motility. **(A)** Laterally-bound kinetochores travel in a directional manner to the plus end of the microtubule, powered by the kinesin-8, Kip3, and the microtubule polymerase, Stu2. In cells, the loss of these alleles results in kinetochore clustering and chromosome congression defects. Together the *in vitro* and *in vivo* data suggests the importance of Stu2 and Kip3 is forming proper end-on kinetochore-microtubule attachments.

## Discussion

For chromosomes to be properly segregated to two daughter cells during mitosis, they must first be aligned at the spindle equator, with each sister chromatid attached to an opposing pole via kinetochore microtubules. Establishment of this configuration through chromosome congression and biorientation, must occur efficiently and accurately to ensure genetic continuity through generations and cellular fitness. Although mitosis has been studied in budding yeast and many other cell types extensively, there are many details of congression that have yet to be elucidated. Due to the compact and crowded nature of the mitotic spindle, it is difficult to visualize individual kinetochore-microtubule attachments using light microscopy, which would be required to describe chromosome congression in detail. The use of a lysate-based reconstitution system from cell-cycle synchronized yeast gave us the unique ability to observe dynamics of mitotic proteins on single microtubules while retaining the complexity of the soluble cellular environment. Consequently, our study was able to shed more light on the mechanism of kinetochore-microtubule attachment and biorientation.

Both Kip3 and Stu2 are required for lateral kinetochore movement on microtubules in metaphase lysates; whereupon reaching the tip, they form end-on attachments (Figure 1). While Kip3 moves much faster than kinetochores in our assay (Figure 2), Stu2 always localizes with the laterally-bound kinetochores, even when Kip3 is absent from the lysate (Figure 3, A and B). Because Stu2 had previously been shown to bind to Kip3, it is a candidate to connect Kip3 to the kinetochore. Adding more credence to the possibility that Kip3 associates with kinetochores is the observation that when six amino acids in the neck of Kip3 (Kip3^NC^), that have been shown to be part of the Stu2-binding region (Gandhi et al., 2011), are mutated to the residues of a mammalian kinesin-8, Kif18a, kinetochores no longer move along the microtubule, even though Kip3 velocity remains unchanged (Figure 3C and D). Of note, Kip3^NC^ does not accumulate on the tips of microtubules and plus-end dynamics shift toward growth and away from depolymerization in *Kip3*^*NC*^ mutants. These observations are consistent with the possibility that these six amino acids might also be important for Kip3’s ability to bind to curved tubulin subunits present at depolymerizing microtubule ends (Arellano-Santoyo et al., 2021; Queen et al., 2023). Another possibility is that Kip3 needs to physically interact with Stu2 at plus ends to accumulate there since loss of Kip3 accumulation at microtubule ends was also observed in *stu2-AID* lysates. In the future it will be important to test these models.

Our evidence indicates that Stu2’s function in kinetochore motility is more complex than simply serving as a link between Kip3 and the kinetochore. While Stu2 depletion from lysates resulted in immobile kinetochores (Figure 1), Kip3 dynamics were also dramatically altered (Figure 3, C and D). While a direct effect of Stu2 on Kip3 motor activity has not been demonstrated, evidence for a direct physical interaction between the two proteins in this and previous studies warrant tests of this possibility (Figure 3, A and B) (Gandhi et al., 2011). Complicating matters further is that the localization patterns for Stu2 and Kip3 on the microtubule are clearly different (Figure 2). One possibility is that Kip3 is stably recruited to kinetochores by Stu2, but the signal is too faint for detection. However, we can conclude that Kip3 motility is directly related to the presence of Stu2 in the extract. When we started with lysates from either strains expressing wild-type Stu2 or depleted for Stu2 using *stu2-AID*, Kip3 motility was instantly altered upon replacement with the other lysate (Supplement Figure2). It is possible that Stu2 is necessary for activating Kip3, or that Stu2 binding alters Kip3 structure in a way that promotes processivity. The possibility that Stu2 binding alters Kip3 conformation is interesting since Stu2’s proposed binding site is in the Kip3 neck-linker G(Gandhi et al., 2011), which has recently been shown to be conformationally important for motility (Arellano-Santoyo et al., 2021; Queen et al., 2023). However, validation of these hypotheses will require further testing to validate. However, since the essential function of Stu2 is tied to its nuclear localization but is independent of its microtubule polymerase activity (Carrier et al., 2022), its function at kinetochores is likely to be important for cell proliferation.

Another layer of complexity is the correlation between impaired Kip3 processivity and immobile kinetochores. Both in *stu2-AID* lysates (Figure 1 and Figure 3) and in wild-type lysates arrested in S-phase and anaphase (Supplement Figure1A), conditions under which Kip3 makes short runs, kinetochores do not move along the microtubule surface. These cell cycle-dependent differences in the motor activity profile of Kip3 could explain why kinetochores are only motile in metaphase lysates (Torvi et al., 2022). We speculate that cell cycle-specific post translational modifications might affect Kip3’s binding to Stu2, the microtubule, or both. Cell cycle specific phosphorylation of Kip3 has been reported (Woodruff et al., 2010; Mieck, 2012). However, site-specific phosphorylation and the direct effects on Kip3’s dynamics have not been studied in this context. It is clear that both Kip3 and Stu2 are significant components in the mechanism of kinetochore motility and are regulated by the cell-cycle. While additional studies are needed to clearly elucidate Kip3’s and Stu2’s exact molecular mechanism, our data provides novel insights into their roles in chromosome congression.

## Methods

### Yeast strains, culturing, and harvesting

Yeast strains used in this study are listed in Table S1. Fluorescent protein fusion tags, degradation tags, and deletion constructs were designed and inserted as described previously (Longtine et al., 1998; Lee et al., 2013; Bindels et al., 2017). Strains were grown in yeast extract peptone dextrose (YPD) rich medium at 25 °C because they contain a temperature-sensitive allele.

Cells were harvested as described previously (Bergman et al., 2018; Torvi et al., 2022). Starter cultures were grown overnight and then diluted into two 2L cultures of YPD at an OD600=6.25×10^−3^. At an OD600=0.4 cultures were shifted to 37 °C for a 3-hour arrest. To specifically degrade a protein of interest, strains containing a degron tag were arrested at 30 °C for 3 hours (total) and then treated with 250 μM 3-indole acetic acid (Sigma-Alrich, St. Louis, MO) for 30 minutes before harvesting. Cells were harvested by centrifugation at 6000 x g in a Sorvall RC5B centrifuge using an SLA-3000 rotor for 10 minutes at 4 °C. Cell pellets were then resuspended in cold ddH2O and pelted in a Hermle Z446K centrifuge for 3 mins at 3430 x g at 4 °C. This last wash step was repeated once more, with any remaining liquid removed from the pellet via aspiration. The cells were then scraped into a 50 mL conical tube filled with liquid nitrogen, snap-freezing them, for storage at −80 °C.

### Generation of whole cell lysates

As described previously (Bergman et al., 2018; Torvi et al., 2022), approximately 5g of frozen cells were weighed into a pre-chilled medium-sized SPEX 6870 freezer mill vial (Spex, Metuchen, NJ). The chamber was then cryo-milled as follows: 3-minute pre-chill, 10 cycles of 3 minutes grinding at 30 impacts per second (15 cps), with 1 minute of rest between grinds. The resulting powered lysate was stored at −80 °C and is usable for 2+ years.

### Generating stabilized, biotinylated, far-red labeled tubulin seeds

As previously described (Bergman et al., 2018; Torvi et al., 2022), purified bovine tubulin was cycled to enrich assembly-competent tubulin. The tubulin was then mixed with both biotin-conjugated and HiLyte 647 (far-red) labeled porcine tubulin (Cytoskeleton Inc., Denver, CO) and resuspended in PEM buffer (80 mM PIPES pH 6.9, 1 mM EGTA, 1 mM MgCl2) to a final concentration of 1.67 mg/mL unlabeled tubulin, 0.33 mg/mL biotin-labeled tubulin, and 0.33 mg/mL far-red labeled tubulin. GMPCPP (Jena Biosciences, Jena, Germany) was added to a final concentration of 1 mM. Aliquots were snapfrozen in liquid nitrogen and stored at −80 °C. Seeds were polymerized immediately prior to use by incubation at 37 °C for 10 minutes.

### Preparation of glass sides and passivation of coverslips

To prepare the glass sides for use in flow chambers, we followed the previously described protocol (Bergman et al., 2018; Torvi et al., 2022). Microscope slides (Corning Inc., Corning, NY) were incubated in acetone for 15 minutes, followed by 100% ethanol for 15 minutes. The slides were then air dried and stored in an airtight container.

To prepare coverslips (1.5 thickness, Corning Inc.) for use in flow chambers, they were first cleaned by sonication in acetone for 30 minutes. Coverslips were then soaked for 15 minutes in 100% ethanol. After three thorough rinses in ddH2O, coverslips were submerged for 2 hours in a 2% Hellmanex III solution (Hellma Analytics, Müllheim, Germany). They were then rinsed in ddH2O three times and dried with nitrogen gas. For passivation, a solution was prepared containing a 0.1 mg/mL mixture of PLL(20)-g[3.5]-PEG(2):PEG(3.4)biotin(50%) (SuSoS AG, Dübendorf, Switzerland), at a ratio of 1:19, in 10 mM HEPES. Coverslips were gently placed on 50 μL drops of this solution on Parafilm and incubated for 1 hour at room temperature in a humid chamber. They were then soaked for 2 minutes in PBS and rinsed in ddH2O for 1 minute. The passivated coverslips were dried with nitrogen gas and stored in an airtight container at 4 °C for a maximum of 3 weeks.

### Assembly of flow chamber

The flow chamber for use in the TIRF-based dynamics assay was prepared and assembled following the procedure described in (Bergman et al., 2018; Torvi et al., 2022).

### Preparation of whole cell lysates for dynamics assay

Similar to the procedure described in (Bergman et al., 2018; Torvi et al., 2022), a 1.5 mL Eppendorf tube was pre-chilled in liquid nitrogen and filled with 0.22 g of powdered lysate. A microtubule length of 5 μm is optimal for visualization. To create microtubules of this length, varying volumes of cold 10X PEM (800 mM PIPES pH 6.9, 10 mM MgCl2, 10 mM EGTA) were added to the lysate. Different strains required different buffer volumes to achieve the desired length in the assay (between 2-15 μL). The lysate was then combined with 0.5 μL of Protease Inhibitor Cocktail IV (Calbiochem, San Diego, CA) and 4 μL of 25.5 U/μL benzonase nuclease (EMD Millipore, San Diego, CA, prod. 70746-3, >90% purity) and thawed on ice for 10 minutes. The cleared lysate supernatant was then transferred to pre-chilled polycarbonate ultracentrifuge tubes and cleared of insoluble material by spinning at 34,600 x g for 25 minutes at 4 °C. Finally, 32 μL of the cleared lysate supernatant was flowed into the previously prepared chamber, as described above.

### Sequential lysate flow assays

The procedures for preparing coverslips, slides, flow chambers, and lysates were followed as described above. However, instead of preparing one lysate per genotype, two lysates were prepared side-by-side. To observe initial behavior, 32 μL of the first lysate was flowed through the chamber and imaged for 7.5 minutes. While the slide was still mounted on the microscope, the movie was paused for 1 minute while the second lysate was carefully flowed in to replace the first. Once the lysate was replaced, the movie was restarted, and the same field of view was captured for another 7.5 minutes. Frames were captured every 2.5 seconds for a total of 15 minutes.

### TIRF Microscopy

After adding the clarified lysate supernatant to the prepared chamber, the slides were loaded onto a Nikon Ti2-E inverted microscope with an Oko Labs environmental chamber prewarmed to 28 °C. Images were collected using a Nikon 60X CFI Apo TIRF objective (NA 1.49) and an Orca Fusion Gen III sCMOS camera (Hamamatsu, Hamamatsu City, Japan) at 1.5X magnification using the Nikon NIS Elements software. A LUNF 4-line laser launch (Nikon Instruments, Melville, NY) and an iLas2 TIRF/FRAP module (Gataca Systems, Massy, France) were used to achieve total internal reflection fluorescence (TIRF), which illuminates the region proximal to the coverslip surface. Images were acquired every 5 seconds for 30 minutes unless otherwise noted.

### *In vivo* live cell microscopy

Cells were grown overnight in YPD medium overnight at 25 °C. The next day, they were diluted into fresh YPD and cultured for approximately four hours (two doublings) at 25 °C until the culture reached log phase (OD600=0.5). The cells were then washed three times with minimal Imaging Medium (synthetic minimal medium supplemented with 20 μg/ml adenine, uracil, L-histadine, and L-methionine; 30 μg/ml L-leucine and L-lysine; and 2% glucose; Sigma-Aldrich) and immobilized on coverslips coated with 0.2 mg/ml concanavalin A. Imaging was performed using a Nikon Ti2-E inverted microscope with an Oko Labs environmental chamber pre-warmed to 25 °C. Images were acquired using a Nikon 60X CFI Apo TIRF objective (NA 1.49) and an Orca Fusion Gen III sCMOS camera (Hamamatsu) at 1.5X magnification using Nikon NIS Elements software. A LUNF 4-line laser launch (Nikon) and an iLas2 TIRF/FRAP module (Gataca Systems) were used for HiLo total internal reflection fluorescence (Tokunaga et al., 2008). Images were taken with 0.2 μm slices for a total 5 μm Z-stack.

### Image and data analysis

For the microtubule dynamics assay, analysis was as described previously (Bergman et al., 2018; Torvi et al., 2022). Imaging data were analyzed using Fiji software (NIH). Registration to correct for stage drift was applied to the raw data (StackReg; (Thévenaz et al., 1998). Kymographs were generated from all microtubules for which the entire length could be tracked for the entire movie. Kymographs were excluded if the microtubules were crossed or bundled. Data from independent technical trials and biological replicates from imaging of one strain were pooled, unless indicated otherwise. For quantification of speeds, a minimum threshold of 86.7 nm/min for kinetochore movement rates was established based on the resolution of the kymographs used to measure speed. Kinetochore movements of less than a threshold slope of 2-pixel displacement (144.4 nm) per 10 pixels of time (50 sec) were categorized as “paused”. Velocities are reported as the mean with the standard error of the mean. Ratios of fluorescent intensities were quantified from individual kymographs that had been background subtracted and bleach corrected using the Fiji plug-in. Using the Fiji “measure” function, average fluorescence intensities were recorded from a line drawn on either the microtubule tip or lattice. Ratios were determined using Microsoft Excel. Statistical significance was determined using a Kruskal-Wallis test (GraphPad Prism, San Diego, CA). P Values are reported as in the figure captions.

For the live cell imaging analysis, images were analyzed using Fiji (NIH). Maximum intensity Z-projections were made, and bleach corrections were applied, using the “histogram matching” macro. Metaphase cells were identified by the presence of a 2-3 μm spindle at the entrance to the bud neck. Cells were then counted as either “bilobed” or “declustered” based on the presence of two distinct kinetochore puncta. Line scans were performed to assist in this binary classification. If two clear peaks were present, the cell was classified as “bilobed.” Any other line scan shape, i.e., 3+ peaks or 1 long kinetochore signal that matched the spindle, was classified as “declustered.” Cells with kinetochores that were classified as “unattached” had at least one kinetochore signal away from and off-axis from the spindle, even though the main kinetochore mass was still “bilobed.” Cells were classified as “both” if there was at least one unattached kinetochore and the main kinetochore mass was also “declustered.” Data are from two independent technical trials of two biological replicates. In each replicate, n = 50 cells were counted. Graphs are of the mean and standard error of the mean (GraphPad Prism).

## Acknowledgements

The authors thank our lab mates, both past and present, for all their support. We also want to thank Matt Miller, Joe Carrier, Erin Jenson, and the whole Miller lab at University of Utah for their Stu2 advice and guidance. This work was supported by the National Institutes of Health (grants R01 GM047842 and R35 GM149237) to G.B. and funds from the Judy Chandler Webb Endowed Chair in the Biological Sciences to G.B. Additionally the work was supported by the University of California Cancer Research Coordinating Committee (CRCC) Predoctoral Fellowship to J.R.T.

## Supplementary Information

**Figure 1.**
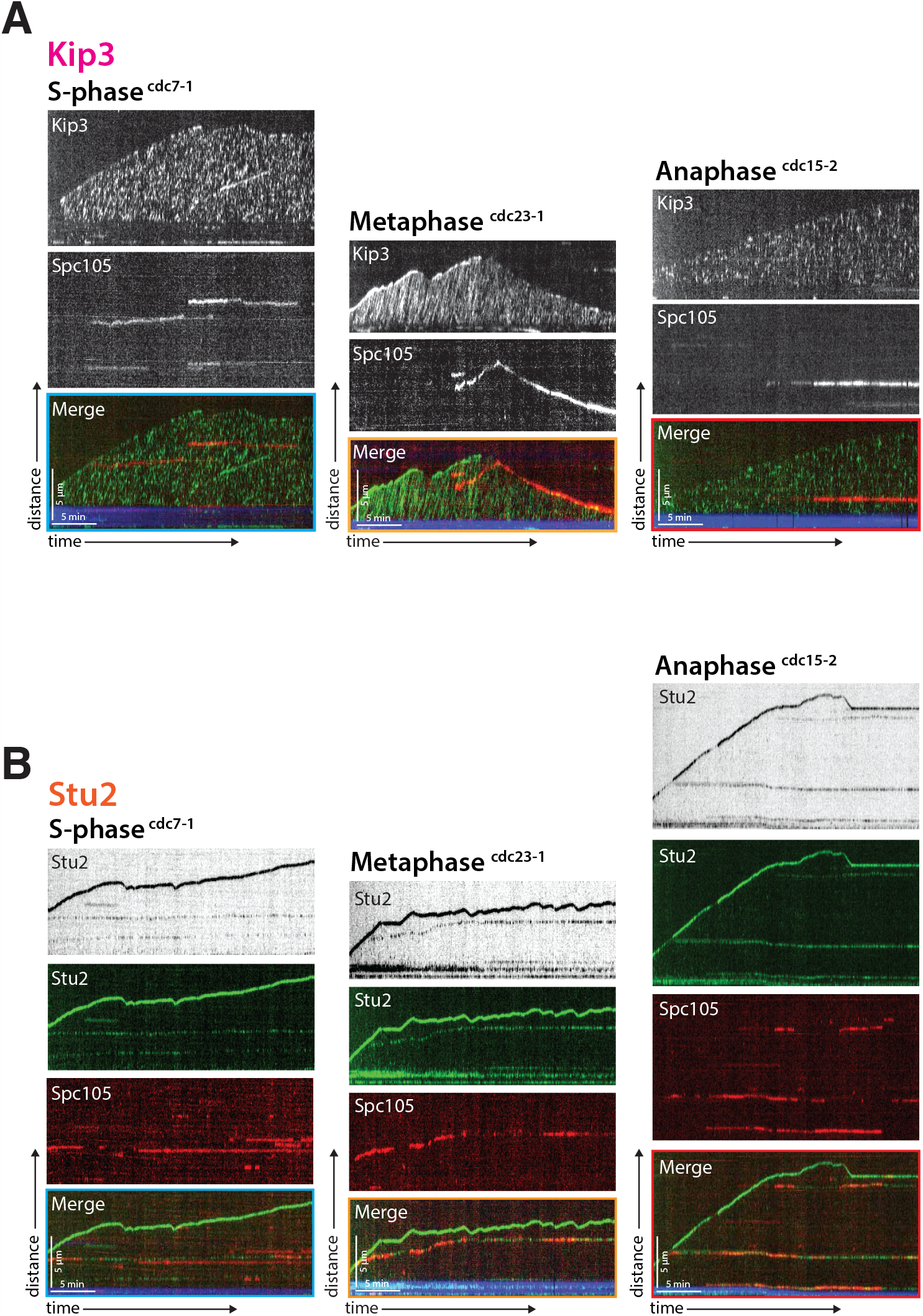
Kip3, Stu2, and the kinetochore dynamics in different cell cycle stages. **(A)** Representative kymographs with Kip3-GFP in green, Spc105-mScarlet in red, and the porcine seed in blue. Time is on the x-axis and distance is on the y-axis (scale bars are 5 min and 5 μm, respectively). Kip3 either comes on and off the microtubule dynamically (S-phase and anaphase) or streams along the microtubule processively (metaphase). **(B)** Representative kymographs with Stu2-GFP in green, Spc105mScarlet in red, and the porcine seed in blue. Time is on the x-axis and distance is on the y-axis (scale bars are 5 min and 5 μm, respectively). Stu2 tracks both the growing and shrinking microtubule tip irrespective of the cell cycle stage. When Stu2 is detected on the lattice, it is invariably colocalized with the kinetochore, marked by Spc105-mScarlet. **(A and B)** Each panel represents different cell cycle arrests using temperature sensitive *cdc* alleles; *cdc7-1* for S-phase, *cdc23-1* for metaphase and *cdc15-2* for anaphase. In metaphase arrested lysates kinetochores, marked by Spc105-mScarlet, are only observed to move toward plus ends, or to track plus ends.

**Figure 2.**
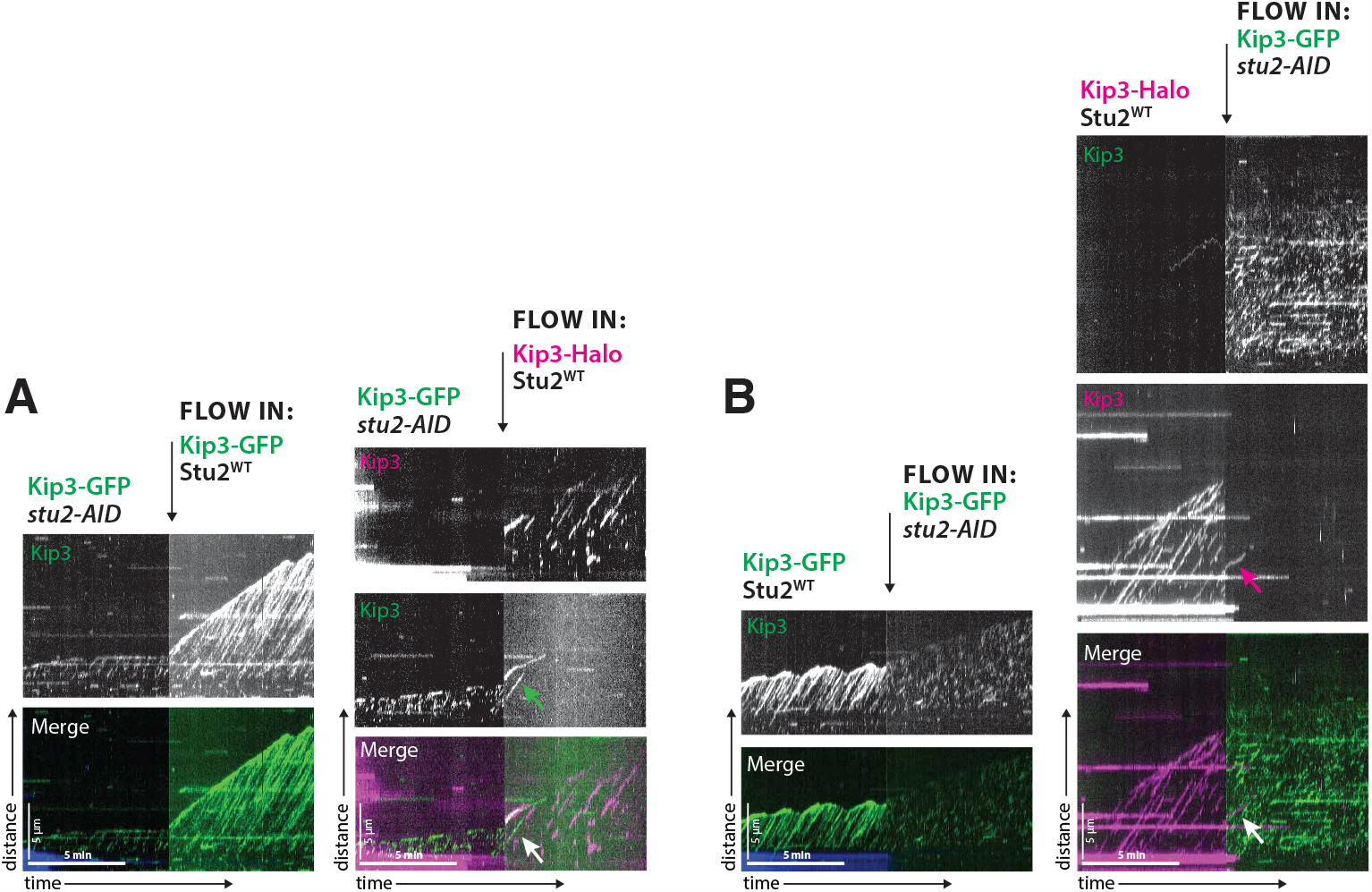
Flow through Stu2 add back to asses Kip3 motility. **(A)** Kip3-GFP was imaged in a *stu2-AID* lysate showing short non-processive motor tracks. Lysate containing wild type Stu2 was added back to the imaging chambers and the Kip3 processivity was tracked either with a Kip3-GFP (left) or with a far red Kip3-Halo (right). Arrows show areas in which Kip3 from the *stu2-AID* strain restored processivity once wild-type Stu2 was added. **(B)** Kip3-GFP (left) or far red Kip3-Halo (right) was imaged in wildtype Stu2 lysate showing long processive motor tracks. Lysate from *stu2-AID* strains, depleted for Stu2, replaced the wild-type lysate in the imaging chambers and the Kip3 processivity was tracked with Kip3-GFP. Arrows show areas in which Kip3 from the wild-type Stu2 strain lost processivity once *stu2-AID* lysate was added in.

**Figure 3.**
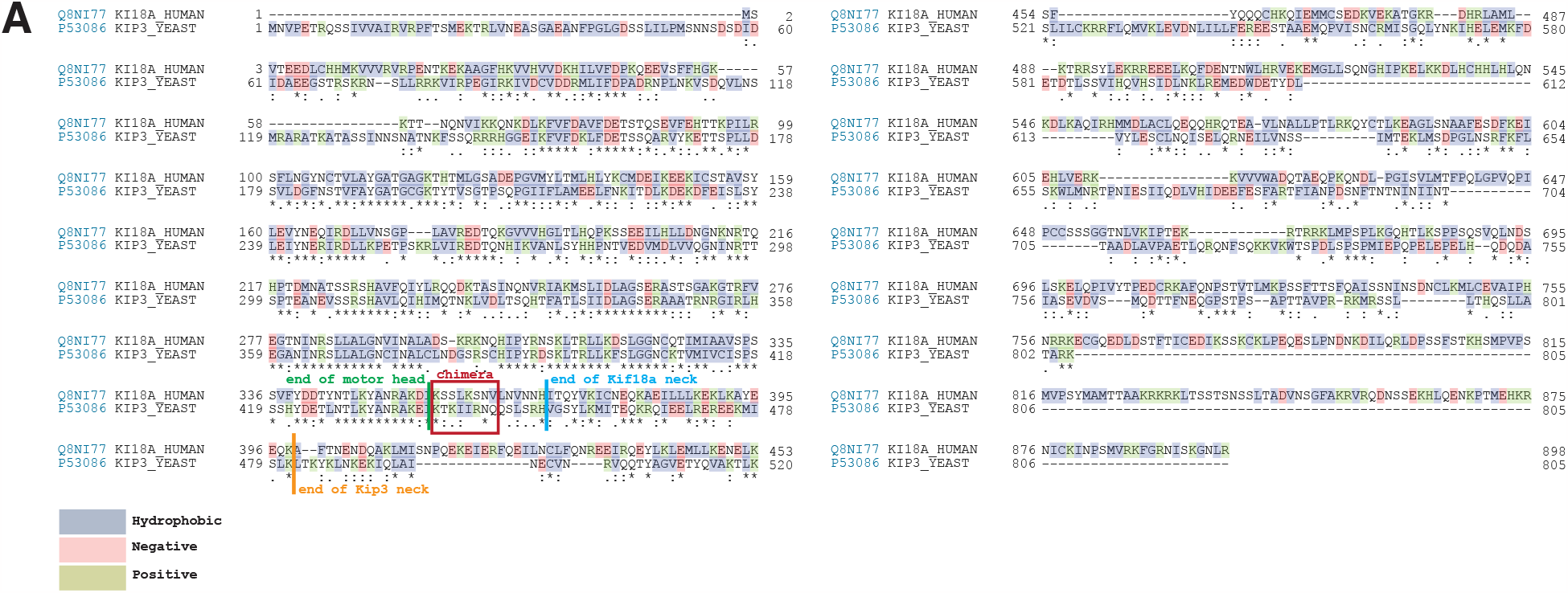
Kip3 neck chimera design. **(A)** The mammalian kinesin-8 sequence was aligned to the yeast kinesin-8, Kip3 using Uniprot.org “align” function. The UniProtIDs are shown on the left, Q8N177 for Kif18a and P53086 for Kip3. Each colored box around an amino acid denotes if it is hydrophobic, negative, or positively charged. A star under the alignment denotes a perfect amino acid match, one circle means they are somewhat similar, and two circles are very similar. Additionally, there is an annotation showing the end of the motor heads, the end of the Kif18a neck, and the end of the Kip3 neck. The red box shows the region of Kip3 that was mutated to match the Kif18a sequence.

## Supplementary Movies

**Figure 1.**
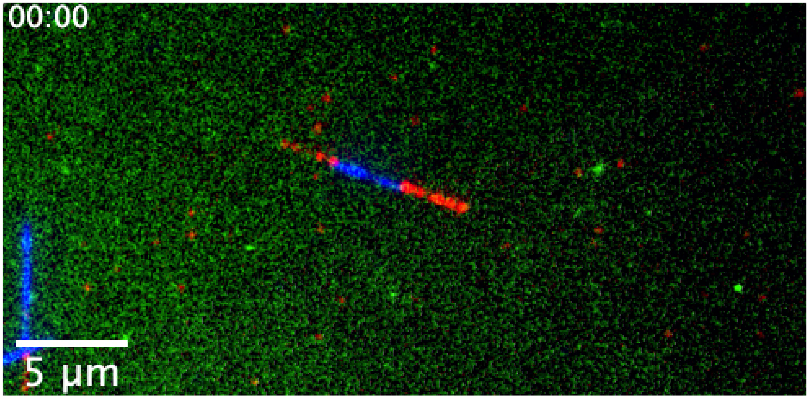
Kinetochore movement on microtubule. Kinetochore identified by Spc105-GFP (green). moves on the lattice of the microtubule (mRuby2-Tub1) (red) toward the plus-end (microtubule seed at minus end, blue). Scale bar is 5 μm. Frame rate is 25 frames per second.

**Figure 2.**
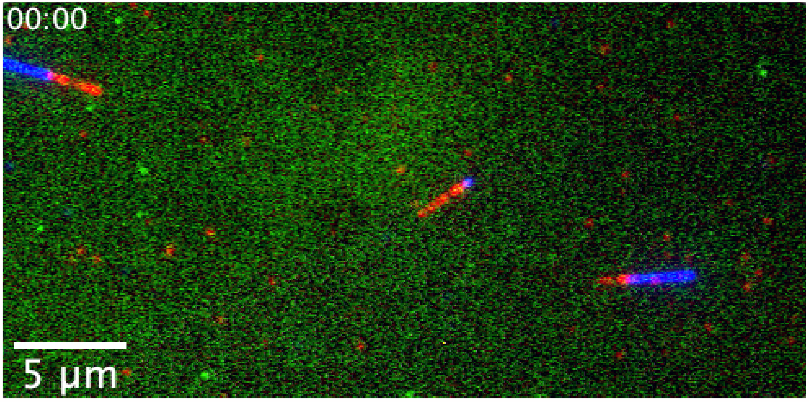
Microtubule-associated kinetochores are stationary when Kip3 is absent. Kinetochores, marked by Spc105-GFP (green), do not move on the lattice of the microtubule (mRuby2-Tub1) (red) toward the plus-end (minus end, blue) when the gene encoding Kip3 is deleted. Scale bar is 5 μm. Frame rate is 25 frames per second.

**Figure 3.**
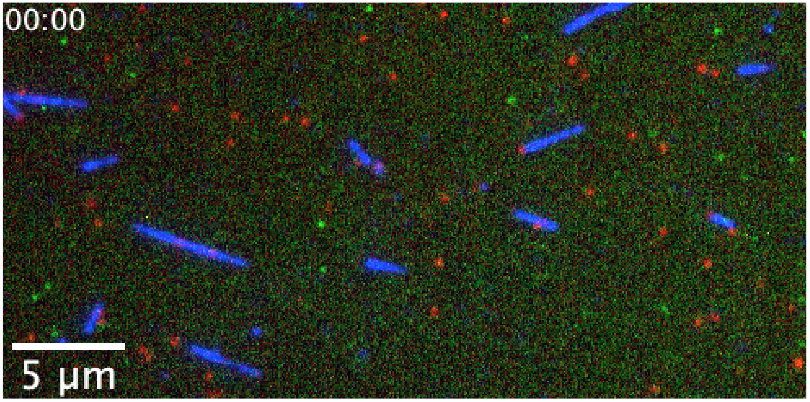
Microtubule-associated kinetochores are stationary when Stu2 is depleted. Kinetochores, marked by Spc105-GFP (green), do not move on the lattice of the microtubule (mRuby2-Tub1) (red) toward the plus-end (microtubule seed at minus end, blue) when Stu2 is depleted using the auxin inducible degron system. Scale bar is 5 μm. Frame rate is 25 frames per second.

**Figure 4.**
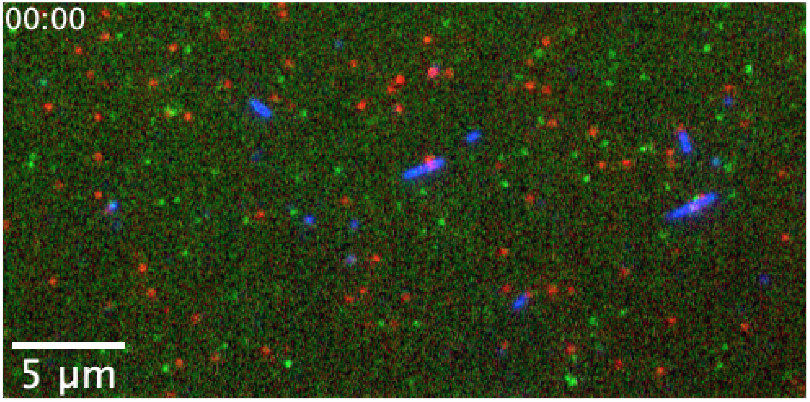
Kinetochores are stationary when both Kip3 and Stu2 are absent. Kinetochores, marked by Spc105-GFP (green), do not move on the lattice of the microtubule (mRuby2-Tub1) (red) toward the plus-end (microtubule seed at minus end, blue) when the gene encoding Kip3 is deleted and Stu2 is depleted. Scale bar is 5 μm. Frame rate is 25 frames per second.

**Figure 5.**
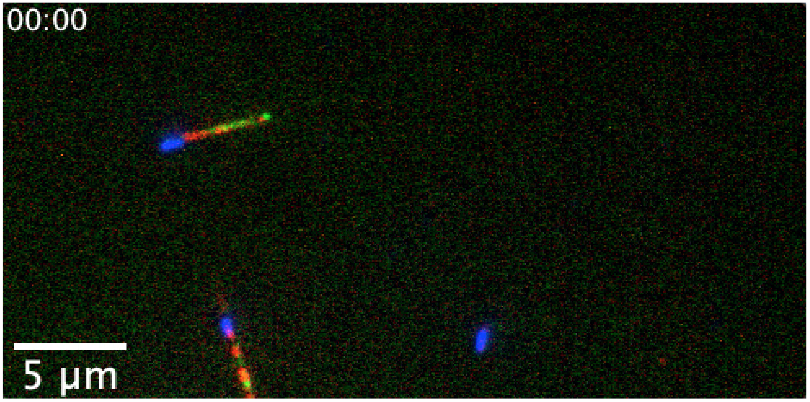
Kip3 dynamics on microtubules. Kip3-GFP (green) walks processively on microtubule (mRuby2-Tub1) (red) toward the plus-end (microtubule seed at minus end, blue). Scale bar is 5 μm. Frame rate is 25 frames per second.

**Figure 6.**
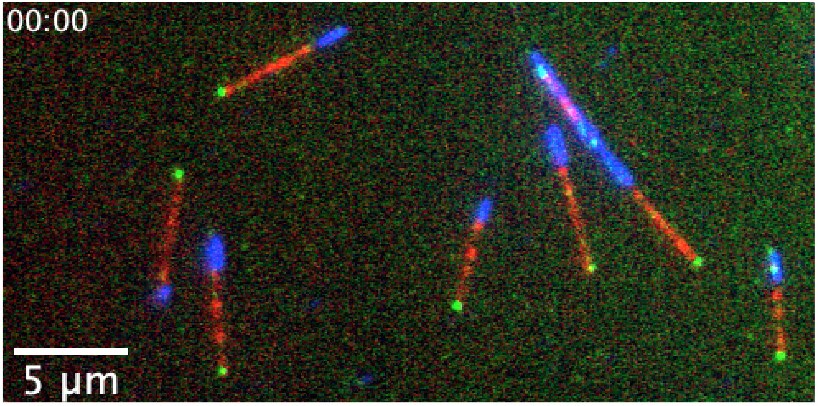
Stu2 dynamics on microtubules. Stu2-GFP (green) tracks the microtubule (mRuby2-Tub1) plus-end (red) but there is also a faint Stu2 signal on the lattice (microtubule seed at minus end, blue). Scale bar is 5 μm. Frame rate is 25 frames per second.

**Figure 7.**
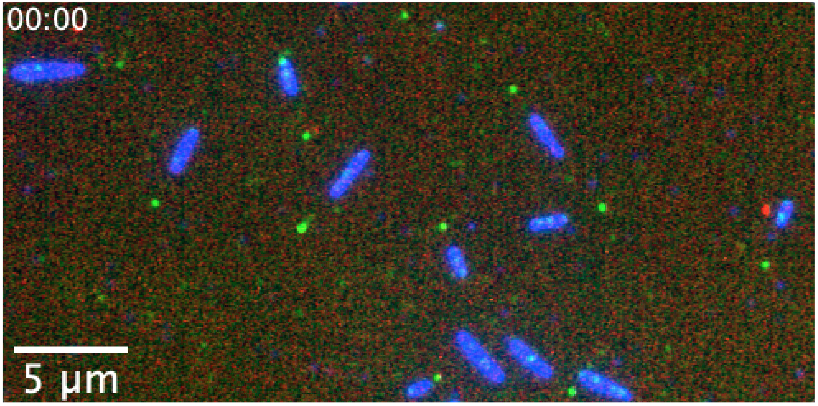
Stu2 and kinetochores colocalize on microtubules. Stu2-GFP (green) when on the microtubule lattice, colocalizes with the kinetochore (Spc105-mScarlet) (red) (microtubule seed at minus end, blue). Scale bar is 5 μm. Frame rate is 25 frames per second.

**Figure 8.**
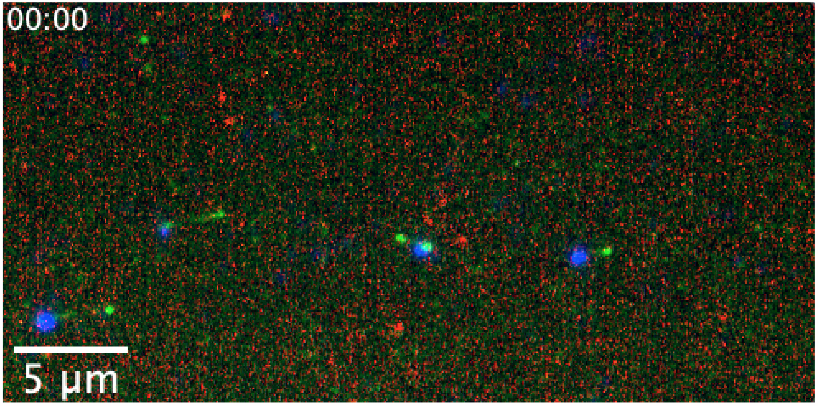
Stu2 and kinetochores still colocalize when Kip3 is absent. Stu2-GFP (green) colocalizes with kinetochores (Spc105-mScarlet) (red), even when the gene encoding Kip3 is deleted (microtubule seed at minus end, blue). Scale bar is 5 μm. Frame rate is 25 frames per second.

**Figure 9.**
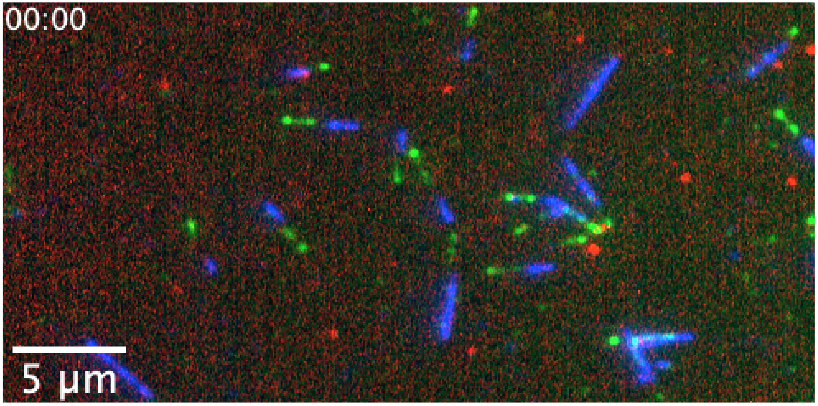
Kip3 and kinetochores do not detectably colocalize. Kip3-GFP (green) moves at a higher rate and is more processive than kinetochores (Spc105-mScarlet) (red), and they do not detectably colocalize (microtubule seed at minus end, blue). Scale bar is 5 μm. Frame rate is 25 frames per second.

**Figure 10.**
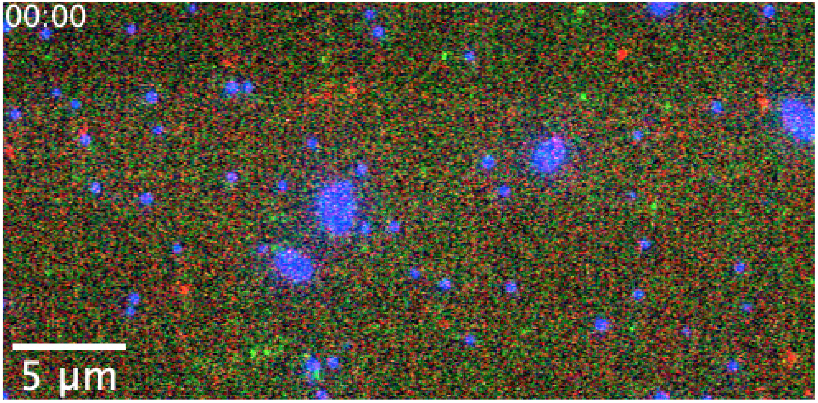
Kip3 processivity decreases but velocity increases when Stu2 is depleted. Kip3-GFP (green) does not colocalize with the kinetochore (Spc105-mScarlet) (red) even though Stu2 depletion alters the motors speed and run length (microtubule seeds at minus end, blue). Scale bar is 5 μm. Frame rate is 25 frames per second.

**Figure 11.**
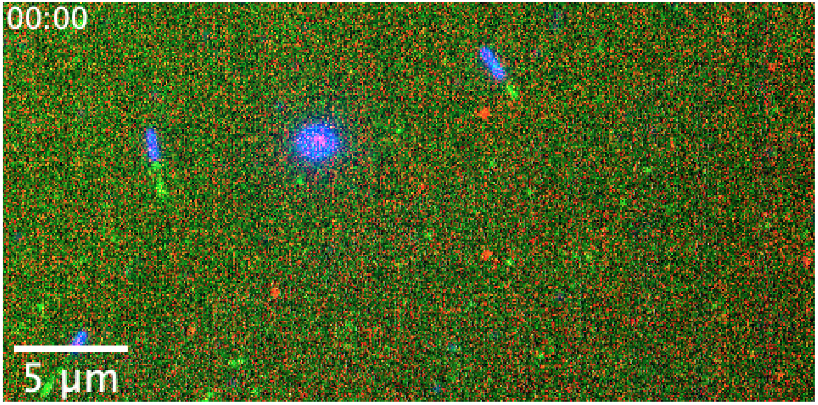
Mutations in the Kip3 neck-linker result in immobile kinetochores but do not affect its own motility. Mutations in the Kip3^NC^-GFP (green) retain motor dynamics but also lead to loss of kinetochore motility (Spc105-mScarlet) (red) (microtubule seed at minus end, blue). Scale bar is 5 μm. Frame rate is 25 frames per second.

## References

Arellano-Santoyo, H., Geyer, E. A., Stokasimov, E., Chen, G. Y., Su, X., Hancock, W., Rice, L. M., and Pellman, D. A tubulin binding switch underlies kip3/kinesin-8 depolymerase activity. Developmental Cell, 42:37–51, 7 2017. doi: 10.1016/j.devcel.2017.06.011.

Arellano-Santoyo, H., Hernandez-Lopez, R. A., Stokasimov, E., Wang, R. Y.-R., Pellman, D., and Leschziner, A. E. Multimodal tubulin binding by the yeast kinesin-8, kip3, underlies its motility and depolymerization. bioRxiv, pages 1–14, 2021.

Ayaz, P., Ye, X., Huddleston, P., Brautigam, C. A., and Rice, L. M. A tog:αβ-tubulin complex structure reveals conformation-based mechanisms for a microtubule polymerase. Science, 337:857–860, 8 2012. doi: 10.1126/science.1221698.

Barisic, M. and Maiato, H. The tubulin code: A navigation system for chromosomes during mitosis. Trends in Cell Biology, 26:766–775, 10 2016. doi: 10.1016/j.tcb.2016.06.001.

Barisic, M., Sousa, R. S. E., Tripathy, S. K., Magiera, M. M., Zaytsev, A. V., Pereira, A. L., Janke, C., Grishchuk, E. L., and Maiato, H. Microtubule detyrosination guides chromosomes during mitosis. Science, 348:799–803, 2015. doi: 10.1126/science.aaa5175.

Bergman, Z. J., Wong, J., Drubin, D. G., and Barnes, G. Microtubule dynamics regulation reconstituted in budding yeast lysates. Journal of Cell Science, 132:1–13, 2018. doi: 10.1242/jcs.219386.

Biggins, S., Gerring, S. L., Connelly, C., Hieter, P., and Dann, P. The composition, functions, and regulation of the budding yeast kinetochore. Genetics, 194:817–46, 8 2013. doi: 10.1534/genetics.112.145276.

Bindels, D. S., Haarbosch, L., Weeren, L. V., Postma, M., Wiese, K. E., Mastop, M., Aumonier, S., Gotthard, G., Royant, A., Hink, M. A., and Gadella, T. W. Mscarlet: A bright monomeric red fluorescent protein for cellular imaging. Nature Methods, 14:53–56, 2017. doi: 10.1038/nmeth.4074.

Carrier, J. S., Torvi, J. R., Jenson, E., Jones, C., Gangadharan, B., Geyer, E. A., Rice, L. M., Lagesse, B., Barnes, G., and Miller, M. P. Stimulating microtubule growth is not the essential function of the microtubule polymerase stu2. BioRxiv, 2022. doi: 10.1101/2022.09.09.507218.

Cottingham, F. R., Gheber, L., Miller, D. L., and Hoyt, M. A. Novel roles for saccharomyces cerevisiae mitotic spindle motors. Journal of Cell Biology, 147:335–349, 10 1999. doi: 10.1083/jcb.147.2.335.

Craske, B. and Welburn, J. P. I. Leaving no-one behind : how cenp-e facilitates chromosome alignment. Essays in Biochemistry, 64:313–324, 2020. doi: https://doi.org/10.1042/EBC20190073.

Gandhi, S. R., Gierlinski, M., Mino, A., Tanaka, K., Kitamura, E., Clayton, L., and Tanaka, T. U. Kinetochore-dependent microtubule rescue ensures their efficient and sustained interactions in early mitosis. Developmental Cell, 21: 920–933, 2011. doi: 10.1016/j.devcel.2011.09.006.

Geyer, E. A., Miller, M. P., Brautigam, C. A., Biggins, S., and Rice, L. M. Design principles of a microtubule polymerase. eLife, 7, 6 2018. doi: 10.7554/eLife.34574.

Heald, R. and Khodjakov, A. Thirty years of search and capture: The complex simplicity of mitotic spindle assembly. Journal of Cell Biology, 211:1103–1111, 2015. doi: 10.1083/jcb.201510015.

Irniger, S., Piatti, S., Michaelis, C., and Nasmyth, K. Genes involved in sister chromatid separation are needed for b-type cyclin proteolysis in budding yeast. Cell, 81:269–277, 1995. doi: 10.1016/0092-8674(95)90337-2.

Kapoor, T. M., Lampson, M. A., Hergert, P., Cameron, L., Cimini, D., Salmon, E. D., McEwen, B. F., and Khodjakov, A. Chromosomes can congress to the metaphase plate before biorientation. Science, 311:388–392, 2006. doi: 10.1126/science.1122142.

Kim, H., Fonseca, C., and Stumpff, J. A unique kinesin-8 surface loop provides specificity for chromosome alignment. Molecular Biology of the Cell, 25: 3319–3329, 11 2014. doi: 10.1091/mbc.E14-06-1132.

Lee, S., Lim, W. A., and Thorn, K. S. Improved blue, green, and red fluorescent protein tagging vectors for s. cerevisiae. PLoS ONE, 8:e67902. 7 2013. doi: 10.1371/journal.pone.0067902.

Lemura, K. and Tanaka, K. Chromokinesin kid and kinetochore kinesin cenpe differentially support chromosome congression without end-on attachment to microtubules. Nature Communications, 6, 2015. doi: 10.1038/ncomms7447.

Li, Y., Yu, W., Liang, Y., and Zhu, X. Kinetochore dynein generates a poleward pulling force to facilitate congression and full chromosome alignment. Cell Research, 17:701–712, 2007. doi: 10.1038/cr.2007.65.

Longtine, M. S., Iii, A. M., Demarini, D. J., Shah, N. G., Wach, A., Brachat, A., Philippsen, P., and Pringle, J. R. Additional modules for versatile and economical pcr-based gene deletion and modification in saccharomyces cerevisiae. Yeast, 14:953–961, 12 1998. doi: 10.1002/(SICI)1097-0061(199807)14:10<953::AID-YEA293>3.0.CO;2-U.

Maiato, H., Gomes, A. M., Sousa, F., and Barisic, M. Mechanisms of chromosome congression during mitosis. Biology, 6:1–56, 2017. doi: 10.3390/|9biology6010013.

Malaby, H. L., Lessard, D. V., Berger, C. L., and Stumpff, J. Kif18a’s neck linker permits navigation of microtubule-bound obstacles within the mitotic spindle. Life Science Alliance, 2, 2019. doi: 10.26508/lsa.201800169.

Mieck, C. Kinesin motor function at the microtubule plus-end, 2012. URL https://core.ac.uk/download/pdf/18263608.pdf.

Miller, M. P., Asbury, C. L., and Biggins, S. A tog protein confers tension sensitivity to kinetochore-microtubule attachments. Cell, 165:1428–1439, 2016. doi: 10.1016/j.cell.2016.04.030.

Miller, M. P., Evans, R. K., Zelter, A., Geyer, E. A., MacCoss, M. J., Rice, L. M., Davis, T. N., Asbury, C. L., and Biggins, S. Kinetochore-associated stu2 promotes chromosome biorientation in vivo. PLoS Genetics, 15, 2019. doi: 10.1371/journal.pgen.1008423.

Queen, K. A., Cario, A., Berger, C. L., and Stumpff, J. Modification of the neck linker of kif18a alters microtubule subpopulation preference. bioRxiv, 5 2023. doi: 10.1101/2023.05.02.539080.

Risteski, P., Jagric, M., Pavin, N., and Tolic, I. M. Biomechanics of chromosome alignment at the spindle midplane. Current Biology, 31:R574–R585, 2021. doi: 10.1016/j.cub.2021.03.082.

Stumpff, J., von Dassow, G., Wagenbach, M., Asbury, C. L., and Wordeman, L. The kinesin-8 motor kif18a suppresses kinetochore movements to control mitotic chromosome alignment. Developmental Cell, 14:252–262, 2 2008. doi: 10.1016/J.DEVCEL.2007.11.014.

Stumpff, J., Du, Y., English, C. A., Maliga, Z., Wagenbach, M., Asbury, C. L., Wordeman, L., and Ohi, R. A tethering mechanism controls the processivity and kinetochore-microtubule plus-end enrichment of the kinesin-8 kif18a. Molecular Cell, 43:764–775, 9 2011. doi: 10.1016/j.molcel.2011.07.022.

Su, X., Qiu, W., Gupta, M. L., Pereira-Leal, J. B., Reck-Peterson, S. L., and Pellman, D. Mechanisms underlying the dual-mode regulation of microtubule dynamics by kip3/kinesin-8. Molecular Cell, 43:751–763, 9 2011. doi: 10.1016/j.molcel.2011.06.027.

Tanaka, T. U., Stark, M. J., and Tanaka, K. Kinetochore capture and biorientation on the mitotic spindle. Nature Reviews Molecular Cell Biology, 6:929–942, 2005. doi: 10.1038/nrm1764.

Thévenaz, P., Ruttimann, U. E., and Unser, M. A pyramid approach to subpixel registration based on intensity. IEEE Transactions on Image Processing, 7: 27–41, 1998. doi: 10.1109/83.650848.

Tokunaga, M., Imamoto, N., and Sakata-Sogawa, K. Highly inclined thin illumination enables clear single-molecule imaging in cells. Nature Methods, 5: 159–161, 2008. doi: 10.1038/NMETH.1171.

Torvi, J. R., Wong, J., Serwas, D., Moayed, A., Drubin, D. G., and Barnes, G. Reconstitution of kinetochore motility and microtubule dynamics reveals a role for a kinesin-8 in establishing end-on attachments. eLife, pages 1–24, 2022. doi: 10.7554/eLife.

Trupinic, M., Kokanovic, B., Ponjavic, I., Barišic, I., Šegvic, S., Ivec, A., and Tolic, I. M. The chirality of the mitotic spindle provides a mechanical response to forces and depends on microtubule motors and augmin. Current Biology, 32: 2480–2493.e6, 6 2022. doi: 10.1016/j.cub.2022.04.035.

Vaughan, K. T. Tip maker and tip marker; eb1 as a master controller of microtubule plus ends, 10 2005. ISSN 00219525.

Woodruff, J. B., Drubin, D. G., and Barnes, G. Mitotic spindle disassembly occurs via distinct subprocesses driven by the anaphase-promoting complex, aurora b kinase, and kinesin-8. Journal of Cell Biology, 191:795–808, 11 2010. doi: 10.1083/jcb.201006028.

Zahm, J. A., Stewart, M. G., Carrier, J. S., Harrison, S. C., and Miller, M. P. Structural basis of stu2 recruitment to yeast kinetochores. pages 1–17, 2021.

